# Muscle Type-Specific Modulation of Autophagy Signaling in Obesity: Effects of Caloric Restriction and Exercise

**DOI:** 10.1101/2024.08.29.610325

**Authors:** Fujue Ji, Jong-Hee Kim

## Abstract

**Background:** Obesity causes metabolic dysregulation and contributes to various diseases, with autophagy playing a pivotal role in this process. Autophagy, a cellular recycling mechanism, is influenced by factors beyond obesity, like caloric restriction (CR) and CR combined with exercise (CR+Ex), which modulate autophagy in obesity management. However, the regulation of autophagy in skeletal muscle under conditions of obesity, CR, and CR+Ex remains poorly understood.

**Method:** Mice were divided into six groups: normal diet, normal diet CR, normal diet CR+Ex, high-fat diet, high-fat diet CR, and high-fat diet CR+Ex. All mice were fed ad libitum with either a normal or high-fat diet for the first four months, followed by the respective interventions for the subsequent four months. Body composition, motor function, and skeletal muscle autophagy signaling were assessed.

**Results:** Obesity resulted in increased total mass, lean mass, fat mass, and fat percentage in tissue; decreased grip strength and endurance capacity. Notably, CR+Ex reduced total mass, lean mass, and fat mass in obese mice. In both normal and obese conditions, the expression of autophagy markers p62, LC3B-I, and LC3B-II is significantly higher in red muscle. Obesity leads to a reduction in cathepsin L expression, while CR further increased LC3B-I expression in red muscle.

**Conclusion:** CR+Ex proves to be an effective strategy for counteracting the adverse changes in body composition associated with obesity. Unlike red muscle, white muscle exhibits lower baseline autophagic levels and may necessitate elevated expression of autophagy-related proteins, such as cathepsin L, to mitigate the negative effects of obesity.

## 1. Introduction

Obesity poses a significant global health challenge, with its prevalence steadily rising across populations [1]. This condition is a major contributor to the development of various chronic diseases, including non-alcoholic fatty liver disease, cancer, diabetes, and chronic kidney disease, largely through its impact on autophagy regulation [2-5]. Understanding the molecular mechanisms underlying autophagy may offer new insights into effective management and treatment strategies for obesity and its related complications.

Autophagy, a fundamental cellular process responsible for maintaining homeostasis, consists of three subtypes: chaperone-mediated autophagy, microautophagy, and macroautophagy [6]. Among these, macroautophagy (hereafter referred to as autophagy) plays a crucial role in the degradation and recycling of damaged cellular components. This process involves the formation of a phagophore, which engulfs cellular debris, formed by its fusion with lysosome to form an autophagolysosome, where degradation occurs [6]. The regulation of autophagy is influenced by both internal and external factors, including obesity, caloric restriction (CR), exercise (Ex), and tissue-specific dynamics [4, 5, 7, 8].

Research has shown that obesity profoundly affect autophagy signaling, leading to functional changes in various organs. In the liver, obesity suppresses autophagy by downregulating key autophagy-related genes such as Atg7, exacerbating insulin signaling and endoplasmic reticulum stress, which contribute to metabolic dysfunction [4]. In contrast, obesity in visceral adipose tissue increases autophagy markers like LC3B and Atg5, which are associated with inflammation and insulin resistance [5]. These disruptions highlight the multifaceted role of autophagy in the pathophysiology of obesity-related metabolic diseases.

Interventions such as caloric restriction (CR) and its combination with exercise (CR+Ex) have demonstrated a significant ability to modulate autophagy and improve metabolic health [7]. CR has been shown to enhance autophagy by upregulating LC3B and Beclin-1, while also improving mitochondrial function [8]. Moreover, CR+Ex further augments autophagic activity, promoting autophagosome formation and enhancing autophagic flux through activating genes such as Atg7, Atg9, and Atg8 [9]. Both CR and CR+Ex inhibit the mTOR pathway, facilitating autophagosome formation and maturation and contributing to the removal of damaged organelles, particularly in the context of obesity and metabolic dysfunction [10].

Beyond their impact on autophagy, CR and CR+Ex have been recognized as non-pharmacological strategies for obesity control [11, 12]. CR, defined by a reduction in calorie intake without malnutrition, has been associated with lifespan extension, reduced body fat, and improved metabolic health, effects partly mediated by enhanced autophagy [13]. The combination of CR and exercise provides even greater metabolic benefits, including improved insulin sensitivity, reduced inflammation, and better lipid metabolism [14]. These interventions not only regulate autophagy directly but also alleviate obesity, establishing a dual mechanism for improving metabolic outcomes [15].

Skeletal muscle, the largest metabolic organ in the body, plays a vital role in energy metabolism,glucose regulation, and fat oxidation, making it integral to overall metabolic health [16]. Autophagy within skeletal muscle fibers is essential for maintaining muscle function by removing damaged organelles and proteins, preventing muscle atrophy, and promoting adaptive muscle regeneration and repair [6, 16]. Given the importance of skeletal muscle in metabolic processes, understanding autophagy in this tissue is critical for addressing muscle function and metabolic disorders related to obesity. Skeletal muscle is composed different muscle types (type I and type II), which exhibit distinct metabolic profiles and responses to stress. Autophagy may play a differential role in these muscle types, influencing their adaptation to various conditions, including obesity, CR, and CR+EX [16, 17]. However, the precise regulatory mechanisms of autophagy in skeletal muscle, particularly under these conditions, remain incompletely understood [16, 18, 19].

This study seeks to advance our understanding of how obesity affects autophagy in skeletal muscle and how CR and CR+Ex modulates these effects across different muscle types. The findings from this study will contribute to the development of targeted therapeutic strategies for obesity-related muscle disorders and provide a scientific basis for designing exercise and nutrition-based interventions.

## 2. Methods

### 2.1 Animals and Experimental Design

Male C57BL/6N mice (n=42, 12-month-old) were randomly assigned to 6 experimental groups: normal diet control (NDC), 20% normal diet calorie restriction (NDR), 20% normal diet calorie restriction + voluntary wheel exercise (NDRE), high-fat diet control (HFC), 20% high-fat diet calorie restriction (HFR), and 20% high-fat diet calorie restriction + voluntary wheel exercise (HFRE).

For the first 4 months, all mice were fed either a standard chow diet (ND; Teklad Global 2018, Envigo Inc.) or a high-fat diet (HF; 45% kcal from fat, D12451, Research Diets Inc.) (Table 1). In the following 4 months, 20% calorie restriction (CR) and 20% calorie restriction + voluntary wheel exercise (CR+Ex) interventions were introduced to the appropriate groups (Figure 1). The mice were housed in a controlled environment, maintained at 22 ± 2 °C with 50–60% humidity, and subjected to a 12-hour light/dark cycle. All animals had ad libitum access to water and were allowed physical activity. The experimental procedures were approved by the Institutional Animal Care and Use Committee (IACUC) of Hanyang University (HYU 2021-0239A).

**Table 1.**
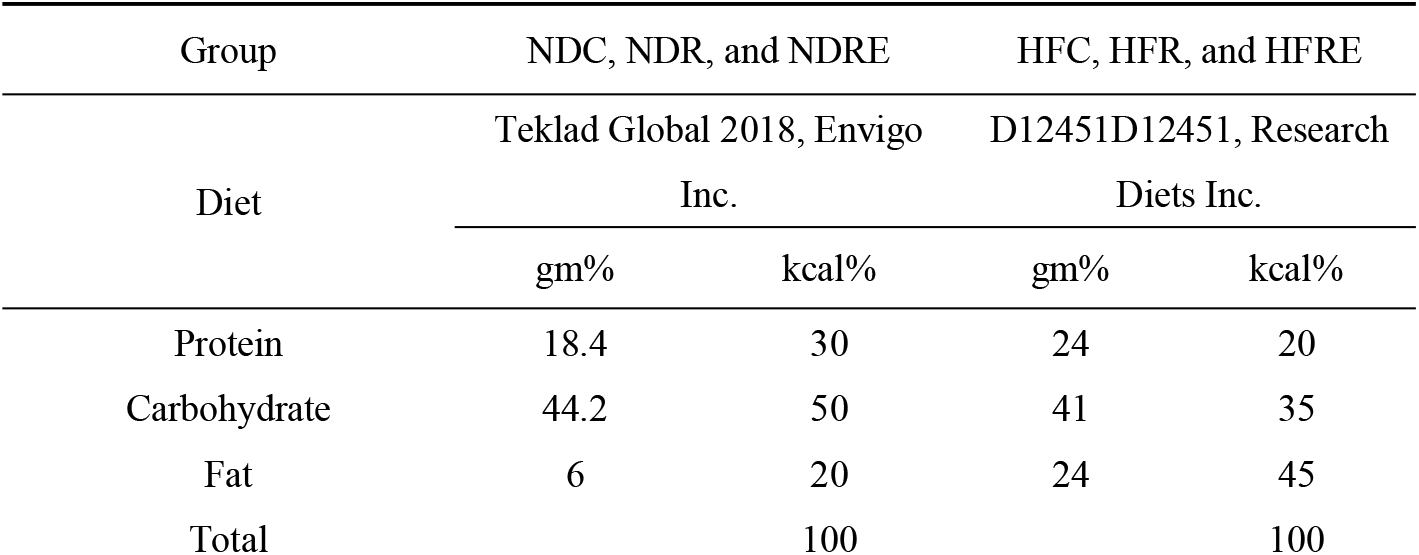
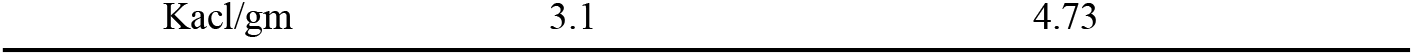
Diet composition.

| Group | NDC, NDR, and NDRE |  | HFC, HFR, and HFRE |  |
| --- | --- | --- | --- | --- |
| Diet | Teklad Global 2018, Envigo Inc. |  | D12451D12451, Research Diets Inc. |  |
|  | gm% | kcal% | gm% | kcal% |
| Protein | 18.4 | 30 | 24 | 20 |
| Carbohydrate | 44.2 | 50 | 41 | 35 |
| Fat | 6 | 20 | 24 | 45 |
| Total |  | 100 |  | 100 |

**Figure 1.**
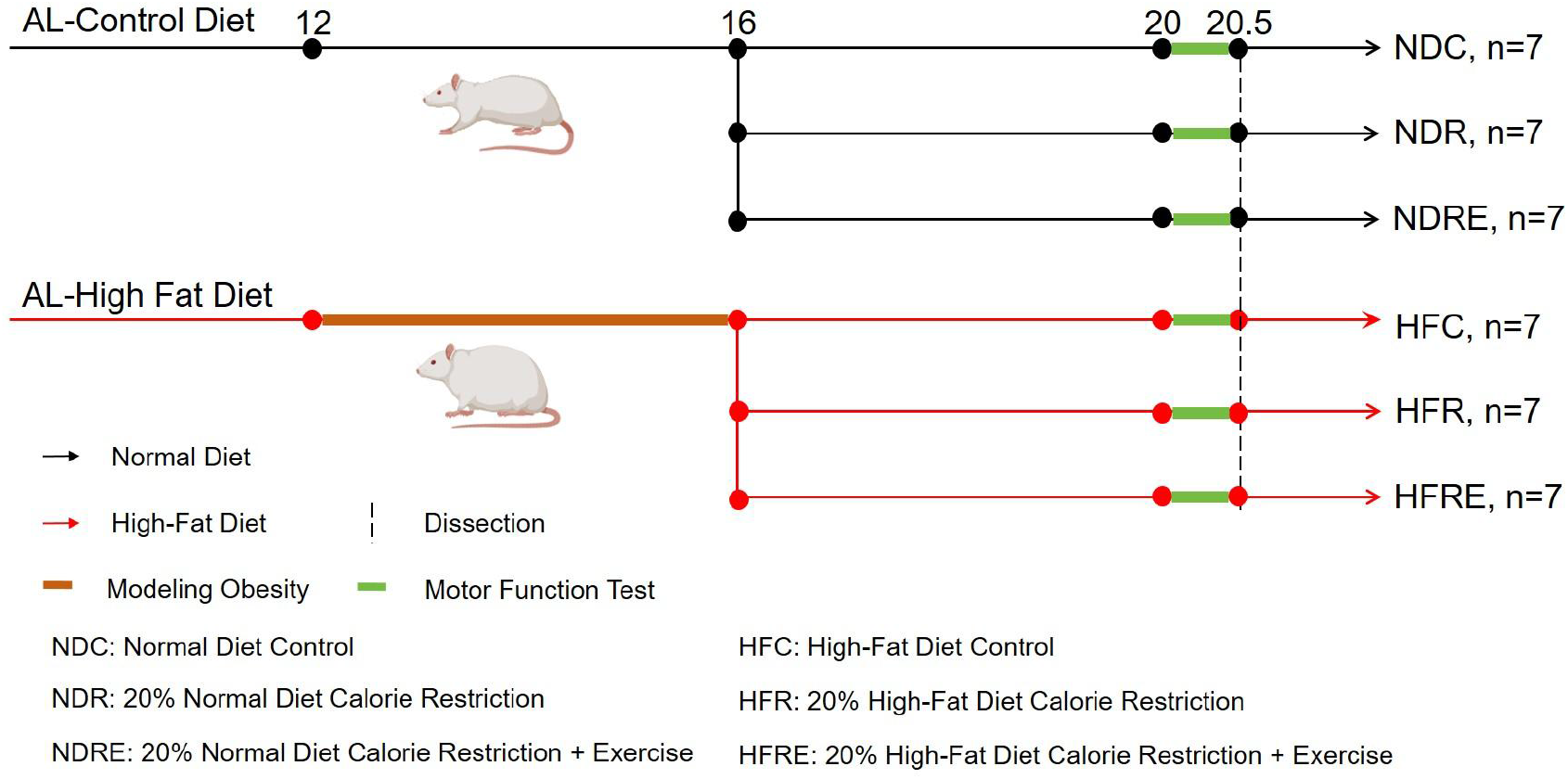
Schematic overview experimental study design.

### 2.2 Motor Function Test

Motor function was assessed in 20-month-old mice two weeks prior to euthanasia. The evaluation included tests to measure walking speed, endurance capacity, physical activity levels, and grip strength across the six experimental groups [20].

#### Walking Speed Test

Walking speed was evaluated using the Rota-Rod test. An adaptation phase was conducted during the first week, in which mice underwent a pre-test session at a constant speed of 5 rpm for 1 minute, once per day. For the official test, the Rota-Rod was set to acceleration mode, gradually increasing the speed from 5 to 50 rpm over a 5-minute period (Model 76-0770, Harvard Apparatus Inc., Holliston, MA, USA). The latency to fall from the device was recorded for each mouse. Each mouse underwent three trials with a ten-minute rest interval between trials, and the best performance among the three trials was used as the final outcome measure.

#### Endurance Capacity Test

Endurance capacity was assessed using a treadmill test. One week prior to the official test, an adaptation phase was conducted in which the mice ran at a speed of 3 cm/s for 10 minutes daily on a 0-degree incline. During the official test, the treadmill speed was increased by 1 cm/s every 20 seconds, starting at 5 cm/s, which the incline set as 0-degrees. The test was terminated when the mouse made contact with the shock pad (set at 0.1 mA) three times.

#### Physical Activity Test

Physical activity was measured using the voluntary wheel running test. Running distance was measured using a voluntary wheel (MAN86130, Lafayette Instrument Company, Lafayette, IN, USA), where each full wheel rotation corresponded to a distance of 0.4 meters. The average running distance over a 5-day period was recorded for each mouse.

#### Grip Strength Test

Grip strength was evaluated using the inverted-cling grip test. An adaptation phase was conducted once daily during the first week. For the official test, each mouse was placed at the center of a wire mesh screen, and a stopwatch was started as the screen was inverted over 2 seconds, positioning the mouse’s head descending first. The screen was held 40-50 cm above a padded surface. The time until the mouse released its grip and fell was recorded. This procedure was repeated three times, with ten-minute intervals between tests, and the longest time recorded was used as the final measurement.

### 2.3 Body Composition

Body composition was examined in 20.5-month-old mice following the motor function test. The mice were anesthetized with 40 mg/kg ketamine and 0.8 mg/kg medetomidine, and body composition analysis was performed using the InAlyzer Dual-energy X-ray Absorptiometry (DEXA) system (Micro Photonics Inc., PA, USA). The measured parameters included total mass (g), fat mass (g), lean mass (g), and the percentage of fat in tissue (%).

### 2.4 Western-Immunoblot (WB)

Following the body composition assessment, the anesthetized 20.5-month-old mice were euthanized via cervical dislocation. After euthanasia, the red and white gastrocnemius muscles were dissected separately [21]. The muscles were lysed using an EzRIPA Lysis Kit (WSE-7420, ATTO), and protein concentration was determined using the Pierce™ Bicinchoninic Acid Protein Assay Kit (Thermo Scientific, Waltham, MA, USA). Equal amounts of protein samples (35 μg) were separated by sodium dodecyl sulfate-polyacrylamide gel electrophoresis and transferred onto nitrocellulose membranes (Bio-Rad Laboratories, Hercules, CA, USA).

The membranes were blocked with 5% non-fat milk dissolved in Tris-buffered saline with Tween-20 (TBST: 10 mM Tris, 150 mM NaCl, and 0.1% Tween-20; pH 7.6) for 1.5 hours at room temperature, followed by overnight incubation with primary antibody (S1) at 4°C. The membranes were then incubated with horseradish peroxidase-conjugated secondary antibody (S1) for 1.5 hours at room temperature. Ponceau S staining was used to normalize protein quantification. The targeted bands were quantified by densitometry using Image J software (National Institutes of Health, Bethesda, MD, USA).

### 2.5 Statistical Analysis

Statistical analyses were performed using GraphPad Prism software (version 9). Two-way analysis of variance with Bonferroni’ s post hoc test was applied. All results are expressed as the mean ± standard deviation (SD). Statistical significance was set at p < 0.05, with asterisks indicating the following levels of significance: *p < 0.05, **p < 0.01, ***p < 0.001, and ****p < 0.0001.

#### 3. Results

### 3.1 CR+Ex improved the changes in total mass, fat mass, and lean mass caused by obesity

Results showed that high-fat diet-induced obesity led to a significant increase in total mass, fat mass, lean mass, and fat percentage in tissue compared to mice on a normal diet (P < 0.05) (Figure 2A-F). Under normal diet conditions, neither CR nor CR+Ex significantly affected body composition (P > 0.05) (Figure 2C-F). However, in high-fat diet-induced obese mice, CR+Ex significantly decreased total mass, lean mass, and fat mass (P < 0.05), although it did not significantly affect fat percentage in tissue (P > 0.05) (Figure 2C-F). These findings suggest that while high-fat diet-induced obesity results in marked alterations in body composition, CR+Ex can improve some of these effects.

**Figure 2.**
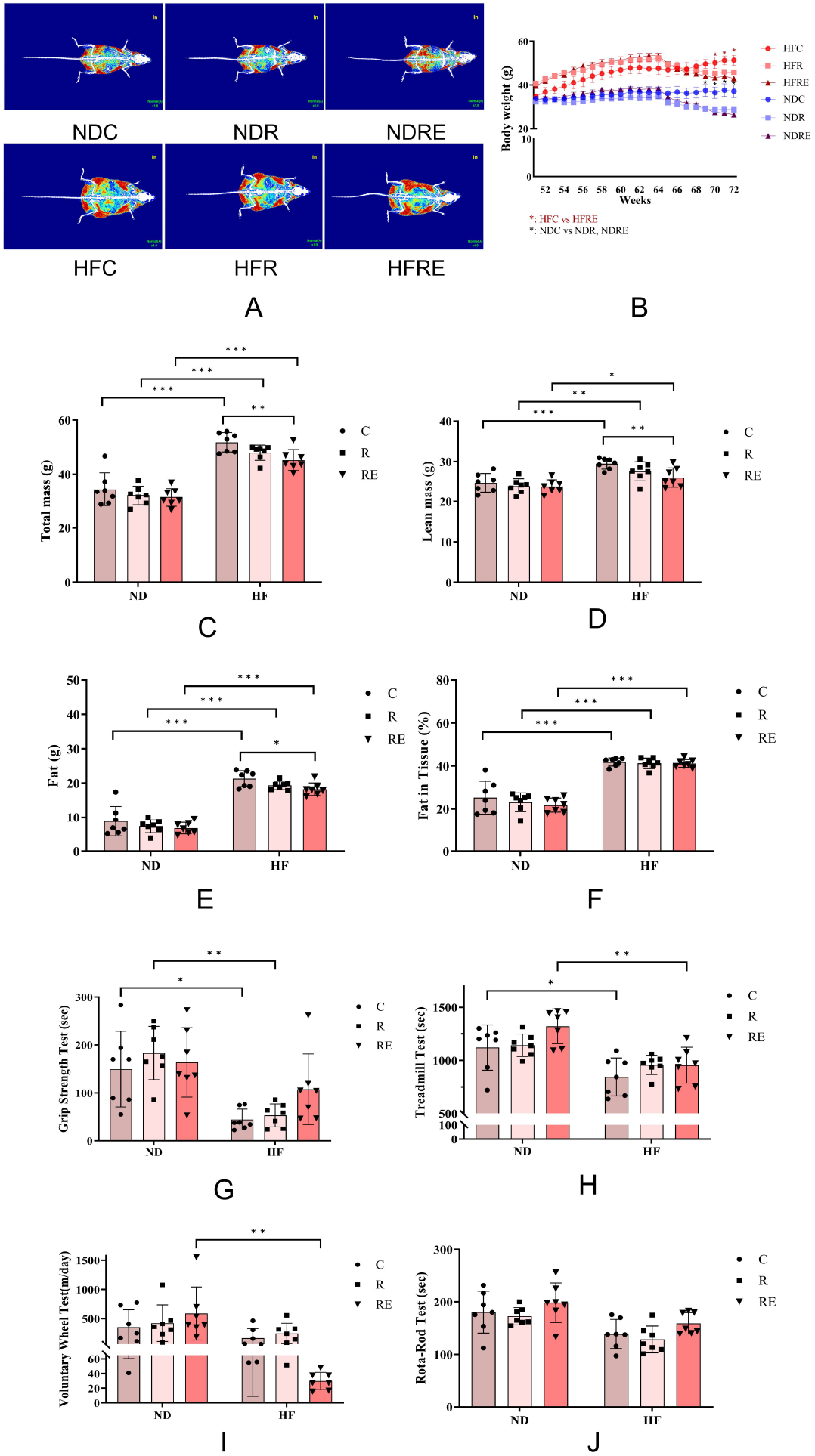
Body composition and motor function results. (A) Representative DEXA-scanned images of NDC, NDR, NDRE, HFC, HFR, and HFRE; skeletal muscle, fat tissue, and bone are displayed in blue, red, and white, respectively. (B) Weekly body weight changes (group × weeks). (C) Total mass. (D) Lean mass. (E) Fat mass. (F) Fat mass in tissue. (G) Grip strength test. (H) Endurance capacity test. (I) Physical activity test. (J) Walking speed test. Significant differences are indicated by asterisks: p < 0.05 (*), p < 0.01 (**), p < 0.001 (***), and p < 0.0001 (****). All values are presented as mean ± SD. Normal diet control (NDC), 20% normal diet calorie restriction (NDR), 20% normal diet calorie restriction + voluntary wheel exercise (NDRE), high-fat diet control (HFC), 20% high-fat diet calorie restriction (HFR), 20% high-fat diet calorie restriction + voluntary wheel exercise (HFRE).

### 3.2 CR and CR+Ex did not rescue the obesity-induced decline in grip strength and endurance capacity

The data revealed that high-fat diet-induced obesity significantly reduced grip strength and endurance capacity compared to a normal diet (P < 0.05) (Figure 2G and H). However, physical activity and walking speed were not significantly affected (P > 0.05) (Figure 2I and J). Notably, neither CR nor CR+Ex significantly improved motor function, regardless of dietary intervention (normal diet and high-fat diet). These findings suggest that CR and CR+Ex were ineffective in reversing the decline in motor function, specifically grip strength and endurance capacity, caused by high-fat diet-induced obesity.

### 3.3 High expression of p62 and LC3B in red muscle indicates active autophagy, independent of interventions

To further investigate the effects of high-fat diet-induced obesity, CR, and CR+Ex on skeletal muscle autophagy, the gastrocnemius muscle was divided into red and white muscle types. First, we examined the interaction between diet (normal diet and high-fat diet) and CR, with or without Ex, on the same muscle type (red or white) to evaluate their specific effects on autophagy singling pathway (Figure 3A). Unexpectedly, the results showed that autophagy related-proteins within the same skeletal muscle type were not significantly affected by the interaction between diet (normal diet and high-fat diet) and CR, with or without Ex (P > 0.05).

**Figure 3.**
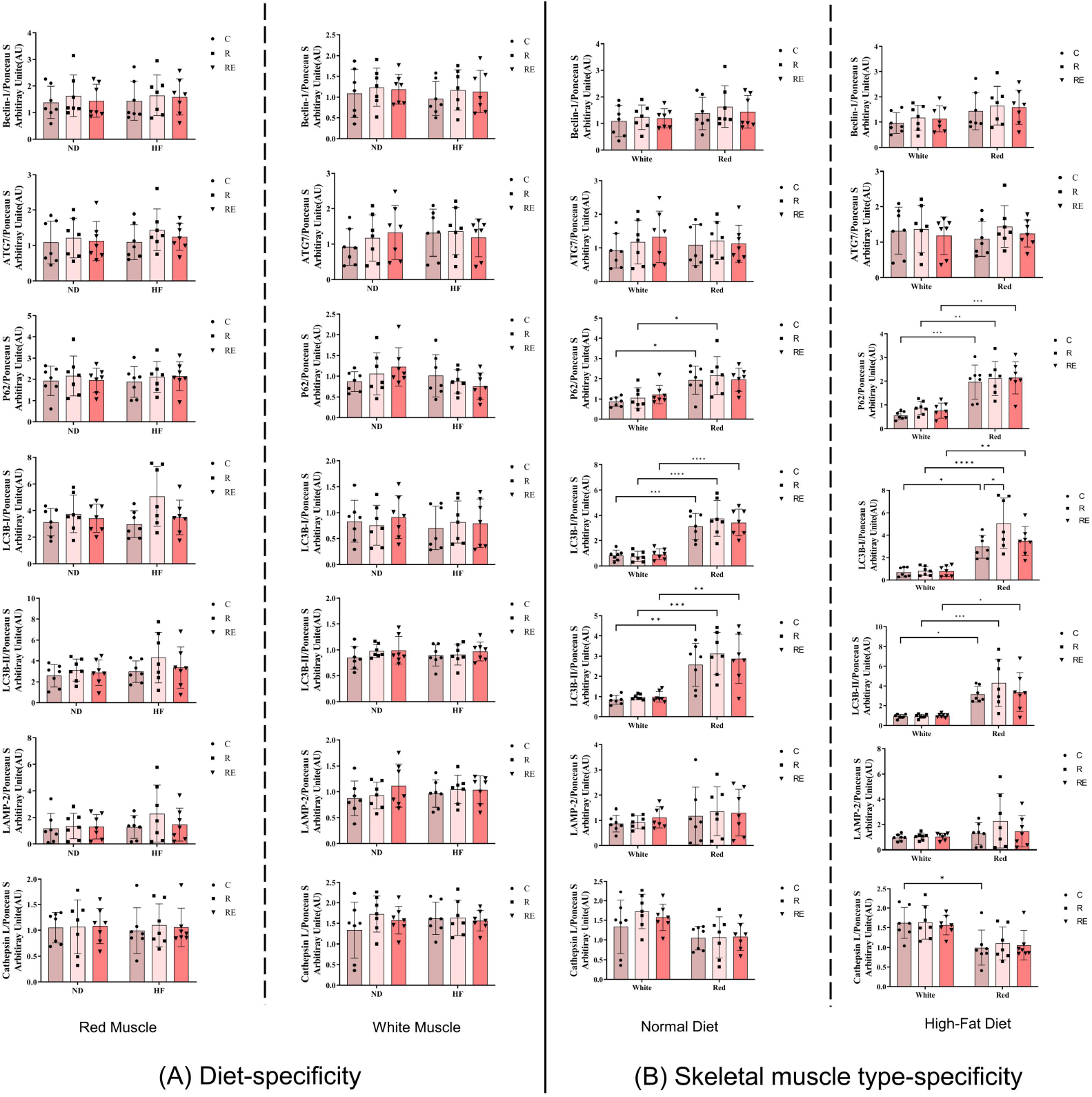
Expression of autophagy-related proteins. Significant differences are denoted by asterisks: p < 0.05 (*), p < 0.01 (**), p < 0.001 (***), and p < 0.0001 (****). All values are presented as mean ± SD. Normal diet control (NDC), 20% normal diet calorie restriction (NDR), 20% normal diet calorie restriction + voluntary wheel exercise (NDRE), high-fat diet control (HFC), 20% high-fat diet calorie restriction (HFR), 20% high-fat diet calorie restriction + voluntary wheel exercise (HFRE).

Next, we examined the interaction between muscle type (red and white) and CR, with or without Ex, under the same dietary conditions (normal diet or high-fat diet) to assess their effects on autophagy singling (Figure 3B). The results showed a significantly higher expression of P62, LC3B-I, and LC3B-II in red muscle compared to white muscle (P < 0.05), regardless of dietary intervention (normal diet or high-fat diet) (Figure 3B). High-fat diet-induced obesity also led to specific alterations in autophagy signaling, particularly with an increased expression of cathepsin L in white muscle compared to red muscle (P < 0.05). Furthermore, under high-fat diet conditions, CR significantly increased LC3B-I expression in red muscle (P < 0.05). These findings suggest that skeletal muscle type plays a more prominent role in the regulation of autophagy-related proteins than dietary interventions, CR, or CR+Ex, with red muscle mostly exhibiting greater autophagic levels than which muscle across different conditions.

## 4. Discussion

Obesity has consistently emerged as a significant topic due to its profound implications for human health [1, 22]. While obesity-induced autophagy dysregulation can contribute to the development of various diseases, there remains a lack of consensus regarding its regulatory mechanisms and potential intervention strategies in skeletal muscle, particularly in light of multiple influencing factors [2-5, 23].

Our experimental findings indicate that the specific regulation of autophagy is predominantly determined by skeletal muscle types rather than by effects of obesity, CR, and CR+Ex. Although obesity and CR can alter autophagy signaling, as evidenced by changes in cathepsin L and LC3B-I levels in red muscle, the regulation of autophagy appears to be significantly more influenced by the inherent characteristics of the skeletal muscle types. These results enhance our understanding of the intricate interplay between skeletal muscle and autophagy mechanisms in the context of obesity, CR, and CR+Ex, providing a scientific foundation for further investigation into cellular degradation mechanisms.

### 4.1 Appropriateness of obesity mouse model

In this study, we employed a high-fat diet to induce obesity in 12-month-old mice, mirroring the gradual increase in obesity rates observed in humans around ages 25-35 [24, 25]. The high-fat diet (D12451, Research Diets Inc.) was administered for four months to establish an obese mouse model, a methodology widely recognized for its effectiveness in inducing obesity [25, 26]. Although there is no universally accepted definition for high-fat diet-induced obesity in mice, most studies implement at least eight weeks of ad libitum feeding on high-fat diet (D12451, Research Diets Inc.) to establish such models [25, 26]. Therefore, a four-month duration of high-fat diet is deemed sufficient to successfully induce an obese mouse model for this study.

### 4.2 Body composition and motor function

Contrary to previous findings [15], we did not observe the anticipated positive effects of CR, likely attributable to differences in CR intensity and the age of the mice in our study. In our study, we employed a lower CR intensity (20%) on older mice (20.5 months), as aging is associated with a slowed metabolism and diminished responsiveness to external stimuli [27]. While high-intensity CR has shown benefits, it may adversely affect older individuals [28]. Consequently, we opted for a lower CR intensity (20%) to ensure the survival of late middle-aged mice and the successful completion of the experiment.

CR+Ex led to alterations in certain aspects of the body composition in the high-fat diet-induced obesity mice, but not in the normal diet group. This discrepancy may be attributed to the higher initial body weight [29] and enhanced fat oxidation efficiency [30], rendering these mice more sensitive to CR+Ex interventions. Overall, obesity induced by a high-fat diet significantly altered the body composition of mice, and CR+Ex positively influenced some of these outcomes.

Our data indicate that high-fat diet-induced obesity leads to reduced grip strength and endurance capacity, consistent with prior study [31]. However, no significant effect on physical activity and walking speed were noted. Surprisingly, neither CR nor CR+Ex significantly enhanced motor function in late middle-aged mice, regardless of dietary interventions. This outcome may stem from the combined effects of lower CR intensity and the specific type of exercise, as well as the reduced responsiveness and adaptability to external stimuli associated with advanced age [27].

### 4.3 Regulation of skeletal muscle autophagy by CR and CR+Ex

In both red and white muscle types, we found no significant effects of the interaction between diet (normal diet and high-fat diet) and CR, with or without EX, on the regulation of autophagy. While such findings are uncommon, similar observations have been reported in the heart [32], skeletal muscle [33], and liver [34]. High-fat diet-induced obesity may disrupt autophagy signaling pathways through various mechanisms. For instance, obesity can impair cellular nutrient-sensing mechanisms, leading to the dysregulated autophagy signaling [35]. Prior research has indicated that, despite the appropriate upregulation of autophagy-related genes at the mRNA level, corresponding protein upregulation may not occur, suggesting a disconnect in autophagy regulation under high-fat diet-induced obese conditions [32]. Additionally, oxidative stress and inflammatory factors, such as ROS, TNF-α, and IL-1β may further inhibit autophagy [36].

In our study, we did not observe significant alterations in autophagy signaling pathways within the same skeletal muscle type due to CR and CR+Ex, regardless of dietary conditions. This is consistent with previous studies, indicating long-term CR and CR+Ex, do not significantly affect the expression of autophagy signaling proteins [7, 10, 32]. This may be related to the age of the mice, the intensity of CR, and the type of exercise used. First, aging weakens external stimuli, reducing the adaptability of the mice [27]. Second, due to the negative effects of high-intensity CR on aging, we used lower-intensity CR to ensure survival rates [28]. However, lower-intensity CR might provide insufficient stimulation for aging mice. Lastly, the voluntary wheel simulates a natural exercise environment for mice, but aging and hunger may reduce the mice’s willingness to exercise, thereby diminishing the effectiveness of the exercise. Although these interventions may upregulate autophagy-related mRNA, this does not consistently translate to notable changes in protein expression level [7, 32].

### 4.4 Regulation of autophagy by skeletal muscle type-specificity

We also examined the differences in autophagy expression between different types of skeletal muscle. Our results demonstrated that P62, LC3B-I, and -II expression levels were significantly higher in red muscle compared to white muscle. Conversely, in high-fat diet-induced obesity, there was a notable increase in cathepsin L expression in white muscle relative to red muscle.

The elevated P62 expression in red muscle compared to white muscle is primarily attributed to differences in metabolic demands and functional roles. Red muscle exhibits greater oxidative metabolic activity, leading to increased ROS production. P62 mitigates ROS-induced oxidative stress by activating the Nrf2 signaling pathway, enhancing the expression of antioxidant enzymes [16, 37]. Additionally, the higher mitochondrial content in red muscle necessitates elevated autophagy to maintain mitochondrial quality, with P62 playing a crucial role in mitophagy [16, 17, 37]. Furthermore, red muscle relies more on fatty acid oxidation, where P62 regulates fatty acid uptake and metabolism, thereby supporting its role in maintaining protein degradation and turnover [16, 17, 37].

Our results show that LC3B-I and LC3B-II expression levels are significantly higher in red muscle than in white muscle, regardless of dietary intervention (normal diet and high-fat diet). This increased autophagic contents in red muscle is essential for maintaining its baseline functions, given its higher oxidative metabolic activity and greater mitochondrial content. Unlike p62, which tags damaged proteins for degradation [37], LC3B-I initiates autophagosome formation, while LC3B-II, the lipidated form, is involved in autophagosome membrane expansion and maturation [38]. LC3B-I and LC3B-II serves as a key marker for monitoring autophagosome formation and flux [38]. Gene and protein expression studies confirm that LC3B-I and LC3B-II levels are significantly elevated in red muscle, reflecting its necessity to meet oxidative metabolic demands and manage oxidative stress, thus ensuring cellular homeostasis [38].

In the context of obesity induced by a high-fat diet, cathepsin L expression was found to be higher in white muscle compared to red muscle, likely due to differences in inflammatory responses, oxidative stress, and autophagic demands [16, 17]. White muscle, which primarily relies on glycolytic metabolism, experiences a heightened metabolic burden when exposed to a high-fat diet, increasing the need for protein degradation and consequently promoting cathepsin L expression [39]. Additionally, chronic inflammation and oxidative stress associated with high-fat diet-induced obesity may render white muscle more susceptible, further elevating cathepsin L levels [40]. Cathepsin L plays a pivotal role in autophagy; thus, the increased demand for protein degradation in white muscle under a high-fat diet conditions drives its elevated expression [16, 17, 40]. In contrast, red muscle, characterized by more active basal autophagy, may not necessitate a significant increase in cathepsin L expression to cope with obesity-induced stress.

In summary, under conditions of obesity, CR, and CR+Ex, autophagy regulation is more pronounced in red muscle compared to white muscle. The differential regulation likely arises from variations in metabolic demands and functional roles, allowing red muscle to adapt more effectively to prolonged low-intensity activities while efficiently executing autophagy to maintain cellular health. Conversely, despite the generally lower basal autophagy levels in white muscle, this tissue compensates for the deficit by upregulating cathepsin L, thereby mitigating the adverse effects associated with high-fat diet-induced obesity.

## Conclusion

CR+Ex represents a viable strategy for ameliorating obesity-induced alterations in body composition. Moreover, the specific expression of autophagy-related signaling in different skeletal muscle types is driven by the necessity to maintain their unique metabolic demands. In particular, red muscle, with its higher baseline autophagic levels, is better equipped to handle external stressors such as obesity. In contrast, white muscle, exhibiting lower autophagic levels, compensates for the deficit by increasing cathepsin L expression. This study provides valuable insights into muscle type-specific autophagy mechanisms in obesity and offers scientific data pertinent to managing obesity and its related health issues.

## Limitation

This study exclusively used male mice, which may not fully represent the responses of other breeds or female mice. Additionally, more advanced research methods, such as spatial proteomics, should be employed to more accurately analyze the muscle type-specific regulation of autophagy pathways.

## Representative Picture of Western-Immunoblot

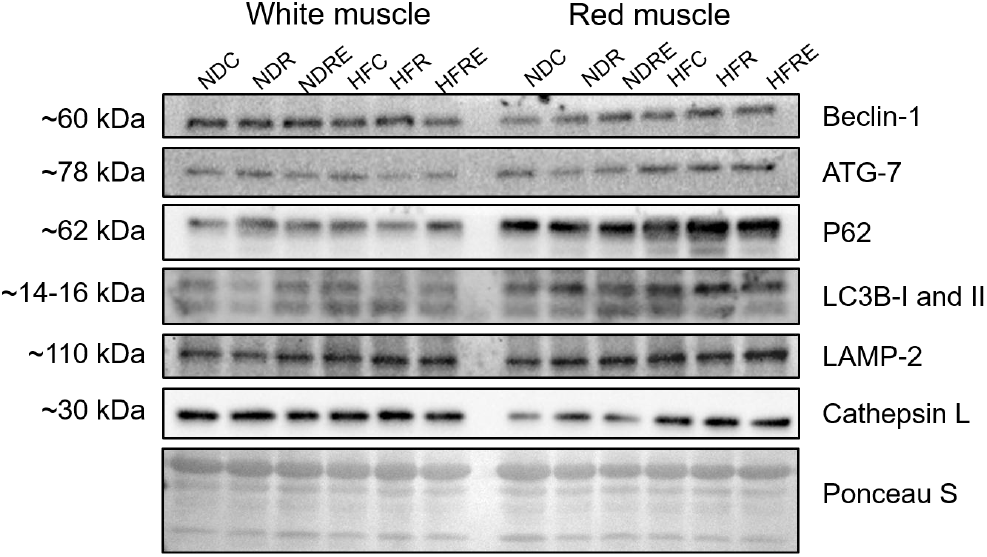

## Consent for publication

Not applicable.

## Availability of data and materials

The data used to support the findings of this study are presented here. Any further data requirements are available from the corresponding author upon request.

## Author’ s contributions

F.J.J. and J.H.K. conceived and designed the study. F.J.J. performed the experiments. F.J.J. analyzed the data and prepared the figures. F.J.J. and J.H.K. interpreted the results, drafted, edited, and revised the manuscript. J.H.K. acquired funding. All authors approved the final version of the manuscript.

## Acknowledgments

We thank Haesung Lee (H.s.L., Hanyang University, Korea) for providing technical support with the statistical analysis of the experimental data.

We thank Jian Guo (Hanyang University, Korea) for his generous help in dissecting the mice. We are grateful to Prof. Dr. Gwang-woong Go (Hanyang University, Korea) for generously providing the DEXA machine for body composition analysis of the mice.

We thank the support of Skill Learning from Kaixin Doctor and MASCU (Medical Association with Science, Creativity, and Unity), Inc, Shenzhen, China.

